# Low risk of transmission of prototype and newly emerged Oropouche virus strains by European *Culex pipiens*, *Aedes albopictus*, and *Anopheles atroparvus* mosquitoes

**DOI:** 10.1101/2025.08.27.672550

**Authors:** Ana Rosales-Rosas, Edith Janssens, Sam Verwimp, Koen Bartholomeeusen, Cédric Van Dun, Katrien Trappeniers, Kevin K. Ariën, Ruth Müller, Leen Delang, Marco Brustolin

## Abstract

Oropouche virus (OROV) is an emerging arbovirus of growing public health concern, with increasing incidence and geographic spread. Since its discovery in 1955, OROV has caused multiple outbreaks in South and Central America, with a new introduction in Cuba since May 2024. Recent travel-related cases in Europe and the Americas underscore its potential for global dissemination. Assessing vector competence outside endemic regions is critical in the context of global travel and climate change.

We evaluated the vector competence of three mosquito species commonly found in Europe—*Culex (Cx.) pipiens, Aedes (Ae.) albopictus,* and *Anopheles (An.) atroparvus*—using two OROV strains: the prototype TRVL9760 (1955, Trinidad and Tobago) and a recent isolate OROV-IRCCS-SCDC_1/2024 (2024, imported from Cuba to Italy). Mosquitoes were orally infected and examined at 7- and 14-days post-infection. We assessed infection (body), dissemination (peripheral tissues), and transmission potential (saliva) by measuring infectious virus particles using the gold standard focus-forming assays.

Our findings show that *Cx. pipiens* and *An. atroparvus* were not susceptible to infection or did not allow transmission with OROV. In *Ae. albopictus*, low infection rates were observed: 6.7% of mosquitoes showed infection at day 7 with the prototype strain, and 3.1% at day 14 with OROV-IRCCS-SCDC_1/2024. All infected mosquitoes showed viral dissemination, but none had infectious virus in their saliva, indicating low risk for transmission.

These results confirm limited vector competence of European mosquito species for OROV and emphasize the importance of continued entomological surveillance to inform future risk assessments.

## Introduction

Oropouche virus (OROV) is an emerging arbovirus and a growing public health concern, marked by its recent increased incidence and geographic expansion. First identified in 1955 in Trinidad and Tobago, OROV was the second most common arboviral disease in South America after dengue, until the emergence of chikungunya (2013) and Zika virus (2015) (1,2). Nevertheless, only nine vector competence studies have been published to date, highlighting the limited research focus of this neglected arbovirus for which no antiviral treatments or vaccines are currently available (3–5).

Traditionally confined to areas near the Amazon rainforest and the Caribbean, OROV propagation and clinical severity have intensified since December 2023. OROV disease is typically mild and self-limiting in immunocompetent individuals with symptoms such as fever, headache, muscle and joint pain, sensitivity to light, and occasionally more severe complications like meningitis or encephalitis. (6,7). However, newly introduced epidemic waves in Cuba since May 2024, along with two fatal cases in healthy Brazilian women, and multiple reports of OROV-linked miscarriages, microcephaly, and fetal deaths, have brought renewed attention to this historically neglected arbovirus (1,8). Additionally, the report of several travel-related OROV cases in countries such as Italy, Spain, Germany, France, the United States, and Canada further underscores the virus’ capacity for international spread (9–11). These developments mark a notable shift in the epidemiological pattern and clinical profile of OROV fever, raising questions on evolutionary changes of the pathogen, the vectors involved in ongoing outbreaks, as well as potential vectors in currently non-endemic areas. Identifying biological adaptations of established versus newly emerged strains is key in understanding altered pathogen behavior. OROV belongs to the Orthobunyavirus genus (Simbu serogroup), characterized by a negative-sense tri-segmented (small [S], medium [M], and large [L]) RNA genome, known to exchange segments with other Simbu strains when co-infecting the same cell. This reassortment drives genetic diversity and viral evolution, leading to potential changes in vector competence and transmission dynamics (12–14). Deiana *et al.* characterized the complete genome of a newly emerged OROV strain (OROV-IRCCS-SCDC_1/2024) isolated from a patient with OROV-induced fever returning from Cuba to Italy in May 2024. This isolate has shown to be a reassortant virus, containing sequences of the Brazil ‘22-‘24 outbreak and older sequences, likely reflecting the strain currently circulating in Cuba and Latin America (15). Whether the observed geographic expansion and disease severity are linked to enhanced virulence in this adapted variant is a current research priority.

Furthermore, a substantial knowledge gap remains regarding our understanding of the reservoirs and vectors involved in OROV transmission (3). Several mammals, including sloths, non-human primates and birds, are considered potential reservoirs for sylvatic OROV maintenance, while no vertebrates other than humans have been identified in urban transmission so far (16). The primary vector of OROV is considered the *Culicoides* midge (family *Ceratopogonidae*), with *C. paraensis* strongly implicated in both sylvatic and urban transmission, given its frequent presence in outbreak zones, which is further supported by experimental data. According to the findings of Pinheiro and colleagues, *C. paraensis* midges are capable of transmitting OROV to hamsters within 6 to 12 days following a blood meal from infected individuals, with the minimum viral titer required for successful transmission and infection of around 5.2 log₁₀ SMLD₅₀/mL (median lethal dose of OROV in suckling mice/mL blood) (17). In addition to biting midges, mosquitoes from the *Culicidae* family could also pose an important vector for OROV transmission. Although OROV has been isolated from various anthropophilic mosquito species since its discovery, the consistently low detection rates and the high experimentally determined threshold of infection (≥9.5 log10 SMLD50/mL), suggested that mosquitoes may not be efficient vectors for human-to-human cycle maintenance (2,3,18–21). However, given the ongoing shifts in vector patterns driven by climate change and global mobility, it is crucial to investigate whether vectors in non-endemic areas can support the transmission cycle of newly emerged OROV strains.

To address this, we evaluated the vector competence of three mosquito species commonly found across Europe: *Ae. albopictus, Cx. pipiens, and An. atroparvus*. These species were exposed to a newly emerged OROV strain (OROV-IRCCS-SCDC_1/2024), isolated from a traveler returning from Cuba to Italy in 2024, or the prototype strain (TRVL9760) originally isolated in 1955. Mosquitoes were examined at 7- and 14-days post infection (dpi) for evidence of infection, dissemination, and transmission potential using the gold standard focus-forming assay (FFA) on Vero E6 cells.

This study expands current knowledge of OROV transmission potential in European mosquito species through the comparison of a historical and an emergent strain. By generating new data on infection, dissemination and transmission, it provides an evidence base to guide surveillance and preparedness in the context of OROV’s recent expansion and its relevance for international public health.

## Material and Methods

### Mosquito species

In this study, three distinct mosquito species were used: *Anopheles atroparvus* (strain Ebre delta, initially collected in Catalonia, Spain 2020) (22), *Culex pipiens* (strain 20CPip.BE-ITM collected in Antwerp, Belgium 2020) (23), and *Aedes albopictus* (strain 21AAlb.IT-TER collected in Terni, Italy, 2021) (24). Established *Aedes* and *Culex* laboratory strains were achieved from the initial field-collected specimens at the Merian Insectarium of the Institute of Tropical Medicine Antwerp (Belgium).

### Mosquito rearing and maintenance

Eggs were placed in trays containing 1.75 L of softened water, supplemented with Novo Fect (Jbl, Neuhoken, Germany) for *Anopheles*-or Koi mini-Sticks (Tetra, Melle, Germany) for *Aedes* and *Culex* larvae development. Colonies were kept in 30 cm × 30 cm × 30 cm cages (BugDorm-1, Megaview, Taichung, Taiwan) with access to 10% glucose solution with 0.1% of Methylparaben (Sigma-Aldrich, Darmstadt, Germany). All mosquito strains were reared and maintained under controlled environmental conditions, 27 °C temperature, 80% relative humidity, in an 11.5/11.5h light/dark cycle with 0.5h of twilight between each cycle.

### OROV stock preparation

The OROV strains used for experimental infections were provided as following: OROV-IRCCS-SCDC_1/2024, a recent isolate from a traveler returning from Cuba to Italy (2024), was kindly provided by Prof. Castilleti (Department of Infectious-Tropical Diseases and Microbiology, IRCCS Sacro Cuore Don Calabria Hospital, Verona), and the prototype strain IP/OROV/TRL9760/12/08/2009, first identified in Trinidad and Tobago (1955) and isolated in France kindly provided by the European Virus Archive Global (EVAg, Marseille, France). Both virus strains were propagated in Vero E6 cells (African Green monkey kidney cells, CRL-1586, ATCC) in Minimum Essential Medium (MEM, Biowest, Nuaillé, France), supplemented with 2% fetal bovine serum (FBS, Sigma-Aldrich, St. Louis, MO, USA), 1% penicillin/streptomycin, 1% L-Glutamine and 1% Sodium Pyruvate (Sigma-Aldrich, St. Louis, MO, USA). In short, cells were infected with virus at a multiplicity of infection (MOI) of 0.01 for 1h at 37°C and 5% CO_2_. Medium was replaced and the culture was further incubated for 24h at 37°C and 5% CO_2_. Supernatant was collected, aliquoted and stored at - 80°C until use. Viral titer was determined by a focus forming assay (FFA) using Vero E6 cells.

### Pairwise distances comparison of the OROV-IRCCS-SCDC_1/2024 strain to the OROV TRVL9760

The sequences for the strains OROV TRVL9760 (accession numbers KC759122-24) and OROV-IRCCS-SCDC_1/2024 (accession numbers PP952117-19) were retrieved from the National Center for Biotechnology Information public database. Using MEGA11 (25), an alignment was built for each OROV segment including both strains, and the pairwise distances were calculated. The percentage of similarity at nucleotide and amino acid level was determined by using the obtained distance value in the formula:

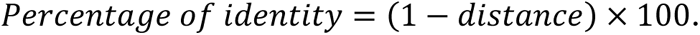

### Vector competence assay

Five-to seven-days-old female mosquitoes were caged in 450 mL cardboard cups (∼60 per species per experiment) and starved for 18h to stimulate feeding. Mosquitoes were exposed to OROV-spiked human blood (final titer of 1 x 10^6^ FFU/mL) via a Hemotek feeding system (SP6W1-3, Hemotek Ltd, Blackburn, UK) through collagen membrane (MEM5, Hemotek Ltd, Blackburn, UK) feeding at 38°C. After 45 minutes of feeding, mosquitoes were anesthetized at 4°C. Fully engorged females were selected on ice, divided in two sub-groups and placed in clean cardboard cups. Mosquitoes were maintained in climatic cupboards under the same controlled environmental conditions used for the rearing with access to cotton soaked in 10% glucose solution. At 7 and 14 dpi, mosquitoes were anesthetized with triethylamine (Sigma-Aldrich, Darmstadt, Germany) prior to mosquito dissection. For each mosquito, only legs (for the OROV-IRCCS-SCDC_1/2024 strain) or legs, wings and head (for the TRVL9760 strain) were dissected first and placed in a 2 ml Eppendorf tube containing 1 mL of mosquito medium (20% FBS in Dulbecco’s phosphate-buffered saline, 50 mg/mL penicillin/streptomycin, 50 mg/mL gentamicin, and 2.5 mg/mL fungizone) containing a zinc-plated steel bead (4.5 mm). Next, mosquitoes were forced to salivate into a 20 µL pipette tip filled with 10 µl of a sucrose:FBS (1:1) solution for 30 minutes, after which saliva was ejected into a 2 mL Eppendorf tube containing 90 µL of mosquito medium. The remaining bodies were placed in 2 mL Eppendorf tubes containing 1 mL of mosquito medium with a zinc-plated steel bead. Body-and peripheral organs-samples were homogenized at 30 Hz for 2 minutes using a TissueLyser II (Qiagen GmbH, Hilden, Germany) and centrifuged for 30 s at 11,000 rpm. All samples were stored at -80 °C until further analysis. Each mosquito species was subjected to two technical replicates for both OROV strains.

### Focus forming assay

To determine vector competence status, the infectious viral load in body, legs and saliva of individual mosquitoes was evaluated by focus forming assay (FFA), as previously described (26–28). In short, samples were serially diluted tenfold and applied to a confluent monolayer of Vero E6 cells, followed by incubation for 24 hours at 37 °C with 5% CO₂ in the presence of 1% carboxymethyl cellulose (CMC) in MEM, supplemented with 5% FBS. After incubation, cells were fixed using 4% paraformaldehyde (Sigma-Aldrich, Darmstadt, Germany) in cold 1× PBS and subsequently blocked with blocking buffer (3% bovine serum albumin and 0.05% Tween-20 in cold 1× PBS). Immunostaining was conducted using a monoclonal antibody against Oropouche virus (Oropouche virus immune ascitic fluid, VR-1228AF, ATCC) diluted 1:1000 in blocking buffer. Following four washes with cold 1× PBS, an Alexa Fluor 488-conjugated goat anti-mouse IgG secondary antibody (Invitrogen, Life Science, Eugene, OR, USA) was used for detection (1:1000 in blocking buffer). Fluorescent foci were visualized and counted using a Leica DMi8 fluorescence microscope (Leica Microsystems, Wetzlar, Germany). Viral titer was expressed as focus-forming units per sample (FFU/sample). Infection and transmission metrics were calculated as follows: the infection rate (IR) was defined as the proportion of mosquitoes with virus-positive bodies among the total number of mosquitoes analyzed following exposure to an infectious bloodmeal; the dissemination rate (DIR) was the proportion of mosquitoes with virus-positive legs (for the OROV-IRCCS-SCDC_1/2024 strain) or virus-positive legs, wings and head (for the TRVL9760 strain) among those with virus-positive bodies; and the transmission efficiency (TE) was the proportion of mosquitoes with virus-positive saliva among the total number of mosquitoes analyzed.

### Statistical analysis

Data analysis was performed using GraphPad Prism version 10.0.2. Differences in infection rate (IR), dissemination rate (DIR), and transmission efficiency (TE) among mosquito species and across time points (7 and 14 dpi). To evaluate differences in viral titers in the body, peripheral organs, and saliva between time points within the same OROV strain, two-tailed Mann-Whitney U tests were employed. Statistical significance was defined as P < 0.05.

## Results

To assess the percentage of similarity of the new isolate to the prototype OROV strain, sequences for both strains corresponding to the S, M, and L segments were aligned respectively, and pairwise comparisons were performed to assess the percentage of divergence at nucleotide and amino acid level. The newly emerged OROV-IRCCS-SCDC_1/2024 isolate (Accession No. PP95117.3, PP95118.3, PP95119.3) was 5.8% and 0% divergent at nucleotide and amino acid level, respectively, in the short (S) segment. The medium (M) segment displayed 5.2% divergence at the nucleotide level and 1.9% at the amino acid level, while the large (L) segment was 10% divergent at the nucleotide level and 5% at the amino acid level (Table 1).

**Table 1.**
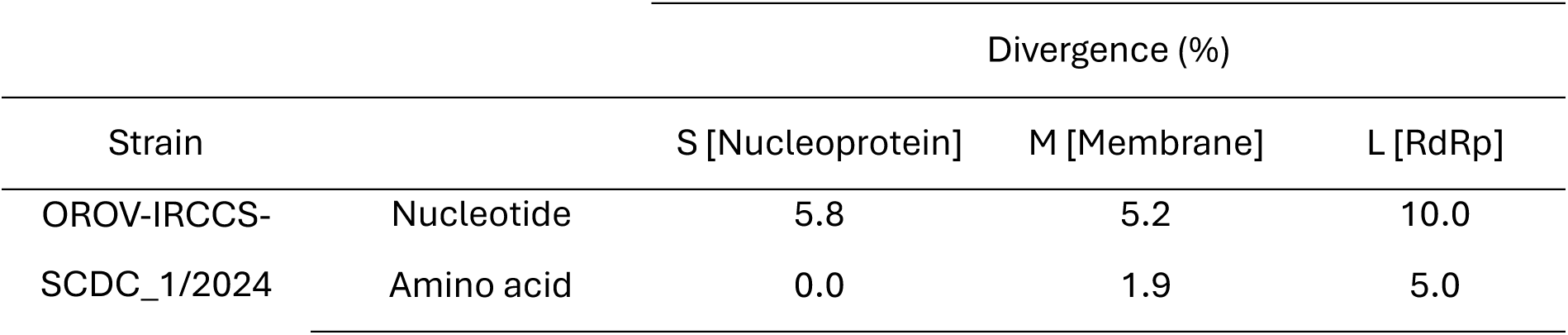
Nucleotide and amino acid sequence pairwise comparisons of the OROV-IRCCS-SCDC_1/2024 strain with the prototype OROV TRVL9760 strain. Segments are presented as S (small), M (medium), and L (large), with the corresponding protein in square brackets. RdRp: RNA-dependent RNA polymerase.

With the prototype OROV strain (TRVL9760), *Ae. albopictus* mosquitoes showed an infection rate of 6.67% (3/45 mosquitoes) at 7 dpi. All three TRVL9760-infected mosquitoes showed a disseminated infection, while no infectious virus could be detected in their saliva (Table 2, Figure 1). The same TRVL9760-infected *Ae. albopictus* mosquitoes had a mean viral titer of 45.6 FFU/sample in the body 7 days post infection, and a mean viral titer of 1.22×10^2^ FFU/sample in the head, wings, and legs (Figure 2). However, no infected mosquitoes could be detected at 14 dpi with this virus strain.

**Table 2.** Experimental infection, dissemination, and transmission outcomes in *Aedes albopictus*, *Culex pipiens*, and *Anopheles atroparvus* mosquitoes following an artificial OROV-infectious blood meal. *IR= Infection rate; DIR= Dissemination rate; TE= Transmission efficiency; n= number of mosquitoes tested*

| Strain | Input PFU/ml | Species | 7 dpi |  |  | 14 dpi |  |  |
| --- | --- | --- | --- | --- | --- | --- | --- | --- |
|  |  |  | IR (n) | DIR (n) | TE (n) | IR (n) | DIR (n) | TE (n) |
| OROV-IRCCS-SCDC_1/2024 | $1 \times 10^6$ | *Ae. albopictus* | 0 (76) | 0 (76) | 0 (76) | **3.12** (64) | **100** (2) | **0** (64) |
|  |  | *Cx. pipiens* | 0 (21) | 0 (21) | 0 (21) | 0 (31) | 0 (31) | 0 (31) |
|  |  | *An. atroparvus* | 0 (22) | 0 (22) | 0 (22) | 0 (19) | 0 (19) | 0 (19) |
| TRVL9760 (prototype) | $1 \times 10^6$ | *Ae. albopictus* | **6.67** (45) | **100** (3) | **0** (45) | 0 (22) | 0 (22) | 0 (22) |
|  |  | *Cx. pipiens* | 0 (40) | 0 (40) | 0 (40) | 0 (42) | 0 (42) | 0 (42) |

On the contrary, infection with the newly emerged OROV strain (OROV-IRCCS-SCDC_1/2024) in *Ae. albopictus* could only be detected at 14 dpi, with an infection rate of 3.12% (2/64 mosquitoes). The two OROV-IRCCS-SCDC_1/2024-infected *Ae. albopictus* mosquitoes showed a disseminated infection, but no transmission of the virus (Table 2, Figure 1). These infected mosquitoes had a mean viral titer of 15.1 FFU/sample in the body at 14 days post infection, and a mean viral titer of 3.15 FFU/sample in the legs (Figure 2). None of the *Cx. pipiens* mosquito bodies or peripheral organs were positive for the TRVL9760 strain, nor for the OROV-IRCCS-SCDC_1/2024 strain at either 7 or 14 dpi. Likewise, *An. atroparvus* exhibited a lack of vector competence for the newly emerged OROV-IRCCS-SCDC_1/2024 strain.

**Figure 1.**
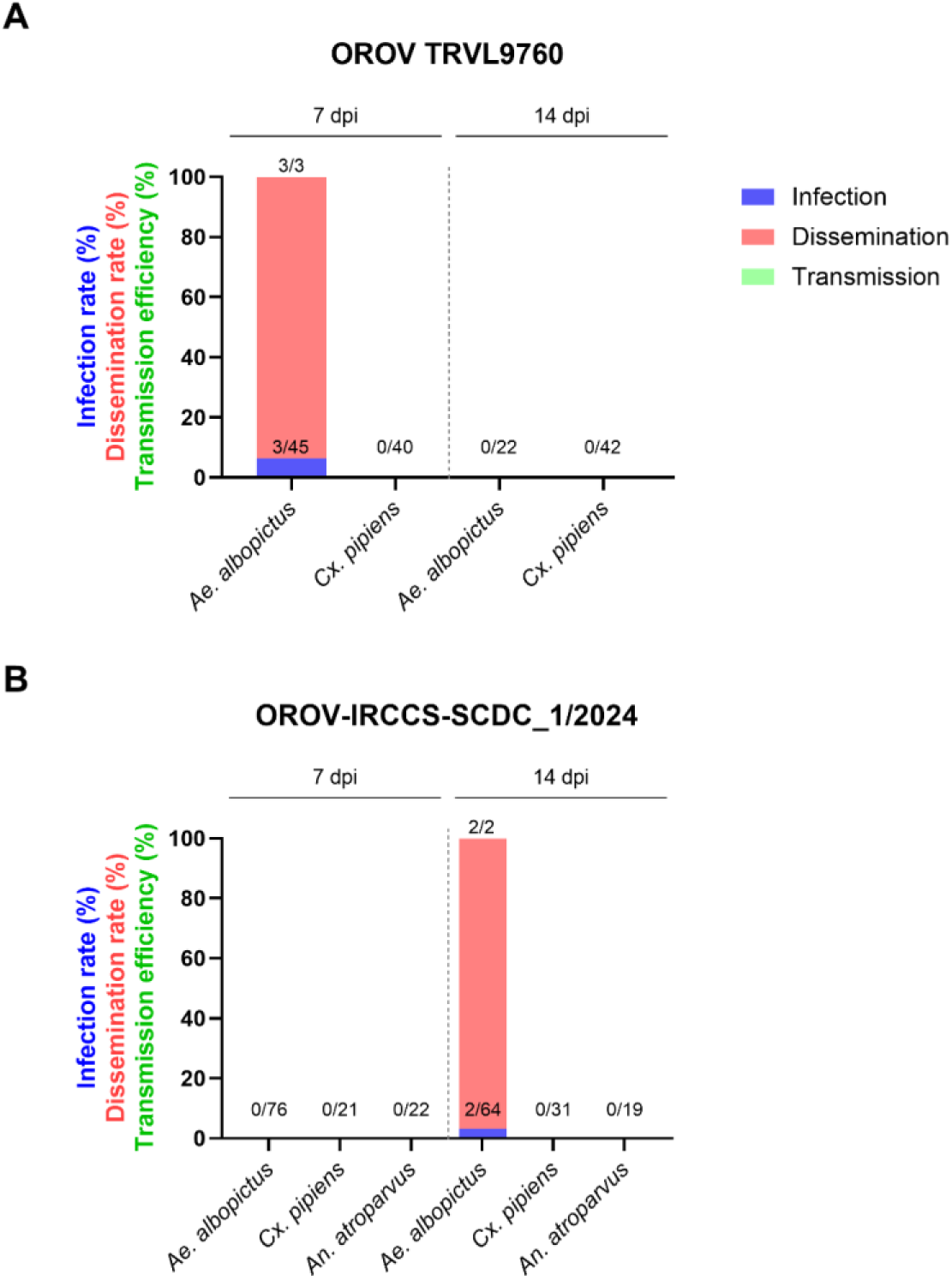
Vector competence of *Culex pipiens*, *Aedes albopictus*, and *Anopheles atroparvus* for OROV strains (A) TRVL9760 and (B) OROV-IRCCS-SCDC_1/2024. The bars indicate the infection rate corresponding to mosquito bodies (blue), dissemination to peripheral organs (red), and transmission potential in the saliva (green) by means of focus forming assay. The labels above the bars represent the number of positive mosquitoes over the total number of mosquitoes

**Figure 2.**
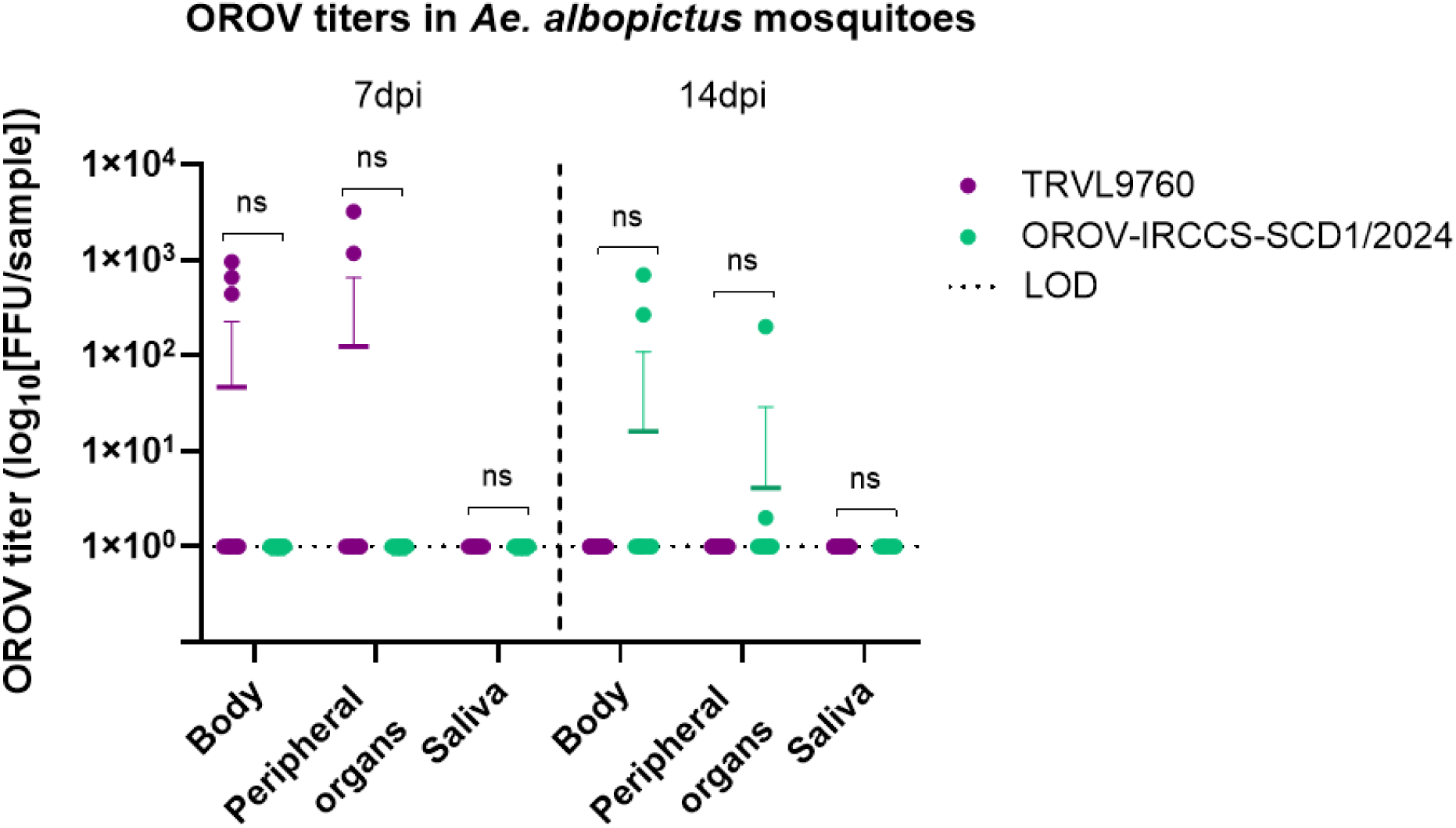
Viral titer (FFU/sample) in the bodies, peripheral organs ((head, wings, and) legs) of *Aedes albopictus* mosquitoes infected with the OROV TRVL-9760 and OROV-IRCCS-SCDC_1/2024 strains at 7 and 14 dpi. For the OROV TRVL9760 strain, head, wings, and legs were included as proxy for dissemination in peripheral organs, while only legs were included for the OROV-IRCCS-SCDC_1/2024 strain. Each symbol represents an individual mosquito sample tested by means of a focus forming assay. The bars represent the mean with SD. To evaluate differences in viral titers in the body, peripheral organs, and saliva between time points within the same OROV strain, two-tailed Mann-Whitney U tests were employed. Statistical significance was defined as P < 0.05.

## Discussion

Our findings demonstrate a low vector competence of *Cx. pipiens*, *Ae. albopictus* and *An. atroparvus* present in Europe for both the prototype and newly emerged OROV strains. Such results are relevant from an epidemiological perspective given their wide dispersal across Southern and more temperate Northern Europe, incl. in Belgium. *Culex pipiens* mosquitoes are native to Belgium and often compose the most abundant species in entomological surveys (29,30). Likewise, *An. atroparvus* is also a Belgian native mosquito; however, its sightings have been scarce in the last nationwide entomological survey carried out (30). Additionally, *Ae. albopictus* mosquitoes have drawn more attention in recent years as an invasive species in Belgium, with dedicated yearly efforts to monitor their presence, as they are known vectors of clinically important arboviruses, including dengue and chikungunya virus (30,31).

Unlike molecular methods detecting viral RNA without indicating infectivity, we employed the gold-standard focus-forming assay (FFA) on Vero E6 cells, directly measuring infectious virus particles, hence providing accurate assessment of the transmission potential.

Our results align with a previous study assessing the vector competence of *Cx. pipiens* mosquitoes from the USA, where no infection, dissemination or transmission of the prototype OROV strain was reported 14 days post infection (32). Nevertheless, these mosquitoes were able to transmit OROV 240023, a strain originally isolated from a febrile patient from Cuba, albeit to a low extent (1 out of 50 mosquitoes). On the contrary, *Cx. pipiens* mosquito populations from the UK were not susceptible to the infection with this OROV 240023 strain (33). Such outcomes hint not only at a strain-specific variability in vector competence, but also at a geographic variation. Furthermore, a recent study reported a lack of vector competence of Italian *Cx. pipiens* mosquitoes (derived from field populations collected in Rome) for the OROV-IRCCS-SCDC_1/2024 strain. These mosquitoes did not show infection or dissemination at 7 nor 14 days post infection (34), which concurs with our findings.

Variations in the kinetics of infection in *Ae. albopictus* mosquitoes were noticed in our study between the employed OROV strains. The mosquitoes that received the prototype OROV strain in the blood meal exhibited infection (6.67%, n= 45) and dissemination (100%, n= 3) already at 7 days post infection, whereas those that received the newly emerged OROV-IRCCS-SCDC_1/2024 strain showed infection (3.12%, n= 64) and dissemination (100%, n= 2) only at 14 days post infection. Payne *et al.* reported a general low vector competence of *Ae. albopictus* (US population) for the prototype OROV strain, with only 1 out of 50 mosquitoes (2%) having a disseminated infection, but no transmission at 14 days post infection (32). On the other hand, Jansen *et al.* have described *Ae. albopictus* mosquitoes (population from Heidelberg, Germany) transmitting the prototype OROV strain at 14 and 21 days post infection, when incubated at both 24°C and 27 °C (35). Our findings regarding the prototype OROV infection kinetics in *Ae. albopictus* differ from what is reported by Payne *et al.* (*32*) and Jansen *et al.* (*35*), as infection and dissemination were detected earlier in our study (7 dpi), and there was no virus transmission. However, the *Ae. albopictus* mosquitoes from our study have originated in Italy; therefore, we should consider that vector competence might be influenced by the possible genetic heterogeneity presented among these mosquito populations, as it has been observed previously for *Ae. aegypti* populations (36).

Concerning the newly emerged OROV-IRCCS-SCDC_1/2024 strain, Mancuso *et al.* found that Italian *Ae. albopictus* (derived from field populations collected in Rome) were infected with OROV when sampled at days 7 and 21 post infection (one infected mosquito out of 20 for each time point); however, no dissemination, nor transmission was detected (37). Likewise, our study employed *Ae. albopictus* mosquitoes derived from an Italian population (collected in Terni) and we did not detect transmission for the newly emerged OROV strain, yet we did observe dissemination at 14 days post infection.

Previous studies have highlighted the need to reconsider the role of *Anopheles* mosquitoes as potential vectors of emerging arboviruses, such as Mayaro virus (38). Payne and colleagues reported a low infection rate (IR = 4%) in An. quadrimaculatus from the United States with Oropouche virus (OROV), with no evidence of viral dissemination or transmission. In our study, we did not detect infection or dissemination in a European population of *An. atroparvus*, suggesting that this species is unlikely to play a role in the transmission cycle of OROV.

Regardless of the time point post infection, the viral titers of OROV in the bodies of *Ae. albopictus* mosquitoes were similar between the two strains (Figure 2). Our study also reports on the amount of infectious virus particles detected in *Ae. albopictus* bodies and peripheral organs following an OROV-infectious blood meal. Other vector competence studies have described that the average OROV viral titer detected in *Cx. tarsalis* bodies 10 days post infection was 31 PFU/mL, whereas the viral titers for *Cx. quinquefasciatus* bodies and leg tissues were 128 and 37 PFU/mL, respectively. At 14 days post infection, *Cx. quinquefasciatus* bodies displayed an average viral titer of 88 PFU/mL, while leg tissues showed an average titer of 100 PFU/mL (39).

All mosquito species tested showed a poor competence for either the prototype TRVL9760 or the OROV-IRCCS-SCDC_1/2024 isolate. When comparing the newly emerged OROV-IRCCS-SCDC_1/2024 strain to the prototype, we found that the highest divergence at the nucleotide level (10%) presented in the L segment, corresponding to the RdRp. This OROV-IRCCS-SCDC_1/2024 strain has been described as a reassortant virus, with the S and L segments sharing high similarity with an emerging cluster of sequences that are most likely related to the recent outbreaks in South America (40). Despite the genetic variations with the prototype OROV strain and the newly emerged strain belonging to a divergent OROV cluster, we did not observe a different outcome in vector competence for the OROV-IRCCS-SCDC_1/2024 strain in our study, for both *Ae. albopictus* and *Cx. pipiens* mosquitoes. Although OROV infectious virions were detected in the bodies and peripheral organs of *Ae. albopictus* mosquitoes, virus was not found in the saliva of any of these mosquitoes, suggesting the presence of a strong barrier to OROV transmission for the tested strains. Nevertheless, the observed viral replication of OROV in certain mosquitoes should not be neglected and underscores the potential for arboviral adaptation and emergence, highlighting the importance of continued surveillance.

Depending on the mosquito-virus combination, typically a higher viral titer in the blood meal can yield a higher percentage of infected mosquitoes (41). While the infectious blood meal in our study contained 1×10⁶ PFU/mL of OROV—a dose within the range typically used in other studies (32,33,42)—we cannot exclude the possibility that a higher viral dose might have resulted in greater viral titers in mosquito bodies and, consequently, altered the transmission potential of Ae. albopictus. Further research assessing the effect of several OROV doses on mosquito vector competence could give a more comprehensive understanding of these infection dynamics. However, the viral input utilized falls on the upper end of the range of viremia detected in OROV-infected humans (6×10^3 and 7×10^5 PFU/ml (43,44)), and therefore, the mosquitoes were exposed to an epidemiologically relevant dose of the virus through the blood meal in this study.

A discrepancy in our study is that dissemination was determined based on different mosquito tissues per strain. The legs were used as a proxy for dissemination for the OROV-IRCCS-SCDC_1/2024 infected mosquitoes, while head, wings, and legs were employed for the prototype TRVL-9760. Consequently, the viral titer measured in the OROV-IRCCS-SCDC_1/2024 infected mosquito legs was lower compared to when using the head, wings, and legs together for the prototype infected mosquitoes (Figure 2). Both tissue types are valid approaches for the assessment of virus dissemination within the mosquito body, as they represent secondary organs that become infected once the viral infection overcomes the midgut barrier. We could observe that the percentage of virus dissemination between both OROV strains remained comparable. As such, this fluctuation in viral titers resulted most likely due to the quantity of tissue tested, and it was further deemed not statistically significant; however, we should remark the low number of mosquitoes that got infected (OROV-IRCCS-SCDC_1/2024, n=2; TRVL-9760, n=3), and thus take this into consideration during comparison.

Similarly to mosquitoes, OROV infection in *Culicoides paraensis* midges has been shown to be dose dependent. However, their threshold for infection is reportedly lower than for mosquitoes, with *C. paraensis* midges exposed to OROV doses higher than 5.2 log_10_ SMLD_50_/mL becoming infected (≥13%) and capable of transmitting the virus (≥40%) (45).

Monitoring of these vector populations in Europe has been triggered during the past decade due to bluetongue and Schmallenberg virus causing considerable economic consequences for European farmers and livestock (46). *Culicoides* midge species are widely distributed across Belgium and Europe with important variation observed in abundance and species diversity between both collection site and sampling period (46,47). While Culicoides species are primarily monitored on farms and focused on identification of species related to infections of food-producing animals, further ecological studies mapping the species’ broader distribution in light of OROV transmission in (peri-)urban settings, is crucial in identifying potential risks of OROV transmission by Culicoides to humans.

While the mosquito species tested in this study appear unlikely to be efficient vectors for both the prototype and a currently circulating OROV strain, ongoing surveillance of both mosquito and *Culicoides* species remains a research priority. Vector- and viral evolution, together with climate change, are expected to alter transmission dynamics, highlighting the value of proactive monitoring to stay ahead of the risks related to the ongoing global spread of this re-emerging virus.

## Ethical statement

The study respects the Directive 2010/63/EU of the European Parliament and of the Council of 22 September 2010 on the protection of animals used for scientific purposes. All animals used are invertebrates (three mosquito species: *Aedes albopictus*, *Anopheles atroparvus,* and *Culex pipiens*). Ethical approval is NOT required.

## Funding statement

This work was supported by the Belgian Directorate General for Development (DGD) (FA5).

The insectaries at ITM are partially funded through the Department of Economy, Science and Innovation (EWI) of the Flemish Government. ARR is supported by a Baekeland Mandate fellowship (HBC.2022.0144) from VLAIO O&O. SV is supported by a PhD fellowship from the Research Foundation – Flanders (FWO) (11D5923N).

## Data availability

All data are fully available.

## Acknowledgements

The authors would like to thank Prof. Joana-Rocha Pereira for kindly providing the prototype OROV stock. We also acknowledged Dr. Concetta Castilletti for providing the OROV-IRCCS-SCDC_1/2024 strain. The authors thank Dr. Nuria Busquets Martí (IRTA-CReSA, Bellaterra, Spain) for providing the authors with the colony of *Anopheles atroparvus* strain Ebre. We thank Maïlis Darmuzey and Martin Ferrié for their advice regarding the foci forming assay for OROV. We also thank our laboratory technicians Jacobus De Witte, Lotte Wauters, Caroline Simons, and Kristien Minner for their help rearing the mosquito colonies. We thank Winston Chiu and Joost Schepers at the Caps-It for their assistance on the imaging and analysis of the foci forming assay plates.

## Conflict of interest

The authors declare there is no conflict of interest

## Contribution

Conceptualization: MB, LD. Methodology: MB, LD, ARR, EJ, KT. Investigation: ARR, EJ, MB, SV, CVD, KT. Formal analysis: ARR, EJ, SV. Visualization: ARR, EJ. Writing – first draft: ARR, EJ, KT. Writing – review and editing: All authors. Funding acquisition: LD, KA, RM. All authors read and approved the final version of the manuscript.

## Notes

### Competing Interest Statement

The authors have declared no competing interest.

